# Characterisation of the butyrate production pathway in probiotic MIYAIRI588 by a combined whole genome-proteome approach

**DOI:** 10.1101/2023.08.20.554021

**Authors:** Liam Wood, Bunmi B Omorotionmwan, Adam M Blanchard, Adam Dowle, Anne L Bishop, Ruth Griffin

## Abstract

Butyrate is a short chain fatty acid with important industrial applications produced by chemical synthesis. With consumer demand for green products, the fermentative production of butyric acid by microorganisms such as *Clostridium* is attracting interest. *Clostridium butyricum* ferments non-digested dietary fibre in the colon to produce butyrate which has multiple health benefits, and certain strains are exploited as probiotics, such as MIYAIRI588 (CBM588). Knowledge of the genes encoding enzymes involved in butyrate production and determining those that are rate-limiting due to low concentrations, could enable strain engineering for higher yields. To this end whole genome sequencing of CBM588 was performed and a circular chromosome, a megaplasmid and the previously reported cryptic plasmid, pCBM588, identified. All genes involved in the butyrate production pathway were found on the chromosome. To identify rate-limiting steps, the relative abundance of the encoded enzymes was assessed by liquid chromatography-mass spectrometry (LC-MS) of total cytosolic proteins. Phosphotransbutyrylase (Ptb) was the least abundant closely followed by butyrate kinase (Buk) and crotonase (Crt). Analysis of upstream regulatory sequences revealed the potential importance of an intact Shine-Dalgarno sequence. Results of this study can now guide bioengineering experiments to improve butyrate yields and enhance the performance of CBM588 as a probiotic.

## 1. Introduction

Butyrate is a four-carbon naturally occurring short-chain fatty acid (SCFA) and has important applications (1). The production of butyric acid is currently through chemical synthesis involving the oxidation of butyraldehyde obtained from crude-oil-derived propylene (2). Approximately 80,000 metric tons are produced annually and sold at around $1.8/kg (3). However, with concerns around the depletion of petroleum resources plus environmental pollution caused by petrochemicals, and consumer preference for bio-based natural ingredients for foods, cosmetics and pharmaceuticals, the production of butyric acid from renewable biomass has become an attractive alternative.

Butyrate is naturally produced in the colon by certain members of the gut microbiota via fermentation of nondigested dietary fibre such as starch (4, 5). Microbiota-derived butyrate is the major energy source for colonic mucosal cells and helps to increase colonic blood flow and increase sodium and water absorption (6). Butyrate is also an important regulator of gene expression for example, it induces colonic regulatory T (Treg) cells which are important for the suppression of inflammatory and allergic responses (7). Butyrate regulates differentiation and apoptosis of host cells resulting in enhancement of the intestinal mucosal immune barrier. Maintaining a certain level of butyric acid production in the lumen not only helps to regulate the host immune response and enhance the intestinal mucosal barrier function but balances the gut microbiota (5, 8).

Butyrate and butyrate derivatives have been used for their therapeutic effects in treating colon cancer, stomach and intestine diseases also hemoglobinopathies (9, 10). Several butyric acid derivatives have been developed as anti-thyroid and vasoconstrictor drugs and used in anaesthetics. In addition to medicinal applications, esters of butyric acid are used as additives to enhance fruit fragrance and as aromatic compounds for production of perfumes (11). Furthermore, in the chemical industry, butyrate is used as a precursor to produce cellulose acetate butyrate (CAB) (12), an important thermoplastic in polymers (13) and paints (14).

The microorganisms that produce butyric acid as the major fermentation product are mainly in the genera of *Clostridium, Butyrivibrio*, and *Butyribacterium* (15, 16). In fermentation using glucose as the carbon source, glycolysis generates pyruvic acid and then acetyl-CoA. Acetyl CoA is subsequently converted either to acetate via the phosphate acetyl transferase (Pta) and acetate kinase (Ack) pathway, or to butyryl-CoA via reactions catalysed by thiolase, acetyl-CoA acetyl transferase (ThlA), hydroxybutyryl-CoA dehydrogenase (Hbd), crotonase/enoyl-CoA hydratase (Crt) and butyryl-CoA dehydrogenase (Bcd). The conversion of crotonyl-CoA to butyryl-CoA by Bcd involves electron-transferring flavoprotein (Bcd-Etfαβ). Butyryl-CoA is then converted to butyrate either by the phosphotransbutyrylase (Ptb) - butyrate kinase (Buk) pathway or via butyryl-CoA:acetate CoA transferase (But). The final step in the acetate biosynthesis pathway and in the butyrate pathway generate energy in the form of ATP (15, 16).

From a biotechnology perspective, *Clostridium* species are the most widely used for the production of butyric acid, and include *butyricum, tyrobutyricum, thermobutyricum, acetobutylicum, beijerinckii*, and *populetii* amongst others (11). The vast majority of butyric acid-producing Clostridia use the Ptb-Buk pathway. However, in *C. tyrobutyricum*, the final conversion of butyrl-CoA to butyric acid is mediated by butyrate:acetate CoA transferase (*cat1*). *C. butyricum* and *C. tyrobutyricum* represent the 2 most promising hosts for butyric acid production due to their strong acidogenic metabolism and ability to tolerate high concentrations of acidic products. *C. tyrobutyricum* is the most studied species with butyrate titres reported to be close to 90 g/L (17). However, this species has a relatively limited substrate spectrum and cannot use polysaccharides or most disaccharides including lactose (11) unlike *C. butyricum* which can produce butyric acid from starch, disaccharides and glucose (18).

*C. butyricum* is found in the environment, for example in soil, and in the intestinal flora of 10–20% of human adults (19). Certain strains have been exploited as probiotics and used in veterinary drugs, feed and food supplements (20). A well-known example of a *C. butyricum* probiotic is strain MIYAIRI588, also known as CBM588 (Miyarisan Pharmaceutical Co., Ltd) which is widely used in Japan, Korea and China to treat antimicrobial induced diarrhoea (21, 22). The bactericidal activities of antibiotics diminishes the normal flora resulting in decreased levels of butyrate-producing bacteria in the colon and impairment of the epithelial barrier, increasing its susceptibility to pathogens (23). CBM588 can counteract gut dysbiosis caused by antibiotics not only by generating SCFAs (butyrate and acetate) but by increasing the abundance of other important SCFAs-producing bacteria, including *Bifidobacterium, Coprococcus*, and *Bacteroides* species (22, 24-28). Additionally, CBM588 has been shown to have antagonistic activity against several enteric pathogens including *Clostridioides difficile, Candida albicans*, enterotoxigenic *Escherichia coli, Salmonella* spp, *Vibrio* spp, *Helicobacter pylori* and enterohaemorrhagic *E. coli* (25-28).

Whilst many microorganisms naturally produce products of industrial or pharmaceutical interest, the rat of production is often hampered by rate-limiting step(s) in the biosynthesis pathway concerned. In order to increase the yields of chosen metabolic products by bioengineering, first it is necessary to identify the main rate-limiting steps in the pathway. Two critical factors that determine the rate of production of metabolites are the concentration of the enzymes involved in the pathway and their level of activity (29). In CBM588, to identify any rate-limiting steps in the butyrate production pathway dictated by the former, first the molecular pathway was determined. Whole genome sequencing of CBM588 was conducted and all genes involved in the butyrate production pathway found based on knowledge of this pathway in another strain of *C. butyricum*, KNU-L09 (30). The products of these genes were then screened by LC-MS of the cytosolic fraction taken from early stationary phase cultures of CBM588 and ranked for abundance.

The data show that Ptb involved in the final stage of the pathway is notably the least abundant enzyme, with levels significantly lower than the most abundant enzyme, ThlA. Ptb was closely followed by Buk and Crt. Intact Shine-Dalgarno (SD) ribosomal binding sites (RBS) were found for all genes involved in the butyrate pathway other than for *ptb, buk* and *crt* which contain single nucleotide polymorphisms (SNPs) suggesting their translation efficiency may be affected. To further investigate whether the presence of an intact SD is associated with stronger expression and conversely SNPs with low expression (as inferred from abundance level), the 40 most abundant proteins were compared with the 40 least abundant. A highly significant difference was found by Mann–Whitney U/Wilcoxon rank-sum test. Our data shed new light on the potential importance of SD RBSs in *C. butyricum* which likely applies to other Clostridial species. With new knowledge from this study and knowledge from previous engineering in *C. tyrobutyicum* (31-33) a set of genetic modifications is proposed for increasing the production of butyrate in CBM588 and potentially enhancing the performance of this probiotic.

## 2. Materials and Methods

### 2.1 Bacterial strain and culture conditions

The commercially available probiotic, *C. butyricum* MIYAIRI588 (CBM588) (Miyarisan Pharmaceutical Co., Ltd) used in this study was first isolated from the faeces of a healthy human by Dr Chikaji Miyairi in 1933, and later from soil in 1963 (21). Unless where stated, CBM588 was cultured in Brain Heart Infusion (BHI) broth or on BHI agar (Oxoid, Basingstoke, UK) supplemented with 0.5% yeast extract (Oxoid, Basingstoke, UK) and 0.1% cysteine (ThermoFisher Scientific, Loughborough, UK) (BHIS) overnight at 37 °C in an anaerobic workstation (Don Whitley Scientific, Bingley, UK) with an atmosphere of CO_2_ (10%), H_2_ (10%) and N_2_ (80%).

### 2.2 DNA extraction and genome sequencing

DNA was extracted from 10 mL culture grown to *A*_600nm_ 0.6 and Illumina sequencing performed as described by Hughes *et al*., (2021) (34). For PacBio sequencing, a DNA library was prepared following the Pacific Biosciences Procedure & Checklist – Preparing whole genome and metagenome libraries using SMRTbell© prep kit 3.0. Two µg of high molecular weight genomic DNA (final volume of 100 µL at 20 ng/µL) were sheared using the Megaruptor3 (Diagenode inc, Denville, New Jersey, USA) at speed The DNA Damage repair, End repair and SMRT bell ligation steps were performed as described in the template preparation protocol with the SMRTbell Express Template Prep Kit 3.0 reagents (Pacific Biosciences, Menlo Park, CA, USA). The sequencing primer was annealed with sequencing primer 3.2 at a final concentration of 1.248 nM and the Sequel II 3.2 polymerase was bound at 0.624 nM. The library went through a SMRTbell beads cleanup (following the SMRTlink v11 calculator procedure) before being sequenced on a PacBio Sequel II instrument at a loading concentration (on-plate) of 120 pM using the adaptive loading protocol, Sequel II Sequencing Kit 2.0, SMRT Cell 8M and 15 hours movies with a 2 hour pre-extension time. The preliminary assembly was performed using the SMRT Link Analysis V11.0.0.146107 pipeline.

### 2.3 Genome assembly and annotation

The PacBio bam file was indexed and converted to fastq using PacBio Tools (v 0.8.1-1) with default parameters. The resulting fastq file was used as input for Flye (v. 2.9.1) (35) using standard parameters with the pacbio-hifi flag. Resulting contigs were assessed for taxonomic assignment using BLAST+ (v 2.13.0) (36). The resulting fasta file was annotated using DFast (v1.2.18) (37) with the reference flag and the reference strain CDC_51208 gff file from accession **<u>GCF_001886875</u>** (38).

### 2.4 Construction of Phylogenetic Tree

Full or scaffold level genomes were downloaded from NCBI in fasta format. Each genome was parsed through SourMash (v 4.5.1) (39) to generate a hash signature which was visualised using Pheatmap (v 1.0.12) (40) in R (v 4.2.1) (R Core Team 2022).

### 2.5 Liquid Chromatography-Mass Spectrometry (LC-MS)

A fresh overnight culture of strain CBM588 was used to inoculate 3 separate 10 mL of *C. difficile* Minimal Medium (41). This medium contains amino acids; 10 g/L Casamino acids, 0.5 g/L L-tryptophan, 0.5 g/L L-cysteine, salts; 0.9 g/L KH_2_PO_4_, 0.9 g/L NaCI, 5 g/L NaHCO_3_, 5 g/L Na_2_HPO_4_, trace salts; 0.001 g/L CoCl_2_.6H_2_O, 0.01 g/L MnCl_2_.4H_2_0, 0.02g/L MgCl_2_.6H_2_0, 0.026 g/L CaCl_2_.2H_2_0, 0.04 g/L (NH_4_)_2_SO_4_, iron; 0.004g/L FeSO_4_.7H_2_O, vitamins; 0.001 g/L D-Biotin, 0.001 g/L calcium D-pantothenate, 0.001 g/L pyridoxine, sugar: 10 g/L glucose.

Cultures were grown at 37°C in an anaerobic workstation to early stationary phase (*A*_600nm_ 2.0). The cultures were centrifuged at 3000 x *g* for 3 min to separate cell pellets from supernatants. To harvest the cytosolic fraction, the pellets were washed with PBS then 5:1 (v:v) Urea Lysis Buffer (20mM HEPES pH 8.0, 9M urea, 1mM sodium orthovanadate, 2.5mM sodium pyrophosphate, 1mM β-glycerophosphate) was added to the cell pellet at room temperature. The slurries were gently pipetted then sonicated at 15W output with 3 bursts of 15 seconds with 1 minute rest between each burst. The lysates were centrifuged at 20,000 x g for 15 minutes at room temperature. Supernatants containing the protein extract were harvested. The protein concentration of the cytosolic fraction was determined by BCA assay (Thermo Fisher Scientific) then frozen at -80°C.

For proteomic analysis, 50 µg of cytosolic fractions were taken for digestion. Protein was denatured in 9M urea before reduction with addition of dithioerythritol and *S*-carbamidomethylated with iodoacetamide prior to diluting to 2M urea with aqueous 50 mM ammonium bicarbonate. Sequencing-grade trypsin/Lys-C mixture (Promega) was added at 1:50 protease to protein ratio before incubation at 37°C for 16 hours. Resulting peptides were desalted with Strata C18-E solid phase extraction (55 μm, 70 Å), dried under vacuum and re-suspended in aqueous 0.1% trifluoroacetic acid (v/v).

Peptides were loaded onto EvoTip Pure tips for desalting and as a disposable trap column for nanoUPLC using an EvoSep One system. The pre-set EvoSep 30 SPD gradient was used with a 15 cm EvoSep C_18_ Endurance column (10 cm x 150 μm x 1.9 μm).

The nanoUPLC system was interfaced to a timsTOF HT mass spectrometer (Bruker) with a CaptiveSpray ionisation source (Source). Positive PASEF-DDA, ESI-MS and MS^2^ spectra were acquired using Compass HyStar software (version 6.2, Bruker). Instrument source settings were: capillary voltage, 1,500 V; dry gas, 3 L/minute; dry temperature; 180°C. Spectra were acquired between *m/z* 100-1,700. Custom TIMS settings were applied as: 1/K0 0.6-1.60 V.s/cm^2^; Ramp time, 166 ms; Ramp rate 5.81 Hz. Data dependant acquisition was performed with 10 PASEF ramps and a total cycle time of 1.89 seconds. An intensity threshold of 1,000 and a target intensity of 20,000 were set with active exclusion applied for 0.4 minutes post precursor selection. Collision energy was interpolated between 20 eV at 0.5 V.s/cm^2^ to 59 eV at 1.6 V.s/cm^2^. High sensitivity detector gain boost was applied.

Peak lists in .raw format were imported into FragPipe (v20.0) (42) for peak picking, database searching and relative quantification. Spectra were searched against the CBM588 chromosome and megaplasmid (GenBank accession numbers CP132344 and CP132345 respectively) appended with common proteomic contaminants and reversed sequences. Search criteria specified: Enzyme, trypsin; Max missed cleavages, 1; Fixed modifications, Carbamidomethyl (C); Variable modifications, Oxidation (M), Acetyl (protein N-term); Peptide tolerance, 12 ppm; MS/MS tolerance, 15 ppm. Peptide identifications were processed using philosopher (v5.0.0) (43) to achieve a 1% false discovery rate assessed against a reverse database. Relative protein quantification was extracted from precursor ion areas using IonQuant (v 1.9.8) (44) with match between runs. MaxLFQ values were used for quantitative comparison post filtering to require protein probabilities of >99.9% and a minimum of two unique peptides per protein.

### 2.6 Statistics

All statistical tests were performed using GraphPad version 7 (San Diego, CA, USA), and *p* values of less than 0.05 were considered to indicate statistical significance.

For analysis of differences between abundance (median of 3 MaxLFQ intensities) for 8 enzymes involved in butyrate production, the data was not normally distributed (determined by Shapiro-Wilk test), therefore the non-parametric Kruskal Wallis test was performed. Post-hoc Dunn’s Multiple Comparisons tests were subsequently performed to determine where the significant differences lie.

To determine whether protein abundance is significantly different for proteins encoded from genes with an intact Shine-Delgarno (SD) sequence compared with non-intact, abundance values (sum of 3 MaxLFQ intensity readings) for the 40 highest and 40 lowest proteins were compared with corresponding SD status using a Mann–Whitney U/Wilcoxon rank-sum test. A non-parametric test was used because the data was not normal (as determined by Shapiro-Wilk test).

## 3. Results

### 3.1 Characterisation of the genome of CBM588

CBM588 was sequenced using a combination of PacBio and Illumina. The long reads generated by PacBio enables resolution of repeat regions. Errors in the long reads are then corrected by short reads generated by the Illumina sequencing. Sequencing and annotation of CBM588 revealed a circular chromosome of 3806640 base pairs harbouring 3349 protein-coding genes, 88 tRNA genes and 30 rRNA genes. Also found was a megaplasmid of 794389 base pairs harbouring 736 protein-coding genes including a CRISPR-Cas operon containing cas6-cas8b-cas7-cas5b-cas3-cas4-cas1b-cas2 (encoded by genes CB_3530 to CB_3600). The chromosome and megaplasmid matched closely to that of *C. butyricum* strain KNU-L09: 98.99% identity, 88% query cover for the chromosome (**<u>CP013252</u>.<u>1</u>**) and 99.22% identity, 76% query cover for the megaplasmid (**<u>CP013489</u>.<u>1</u>**) upon BLASTn analysis in NCBI. The previously identified cryptic plasmid of CBM588, pCBM588, (**<u>AB365348)</u>** harbouring 9 protein-coding genes including a bacteriocin (ORF3) (45) was additionally identified as expected. The GenBank accession numbers for the chromosome, megaplasmid and cryptic plasmid deposited with the NCBI are **<u>CP132344</u>, <u>CP132345</u>** and **<u>CP132346</u>** respectively.

### 3.2 Phylogenetic analysis of CBM588

CBM588 was compared to other complete or scaffold genomes which are publically avalible on NCBI and CDC_51208 used as the reference strain (38). CBM588 clustered with two other strains (TOA and KNU-L09) and also closely clustered to strain JKY6D1 (Figure 1).

**Figure 1.**
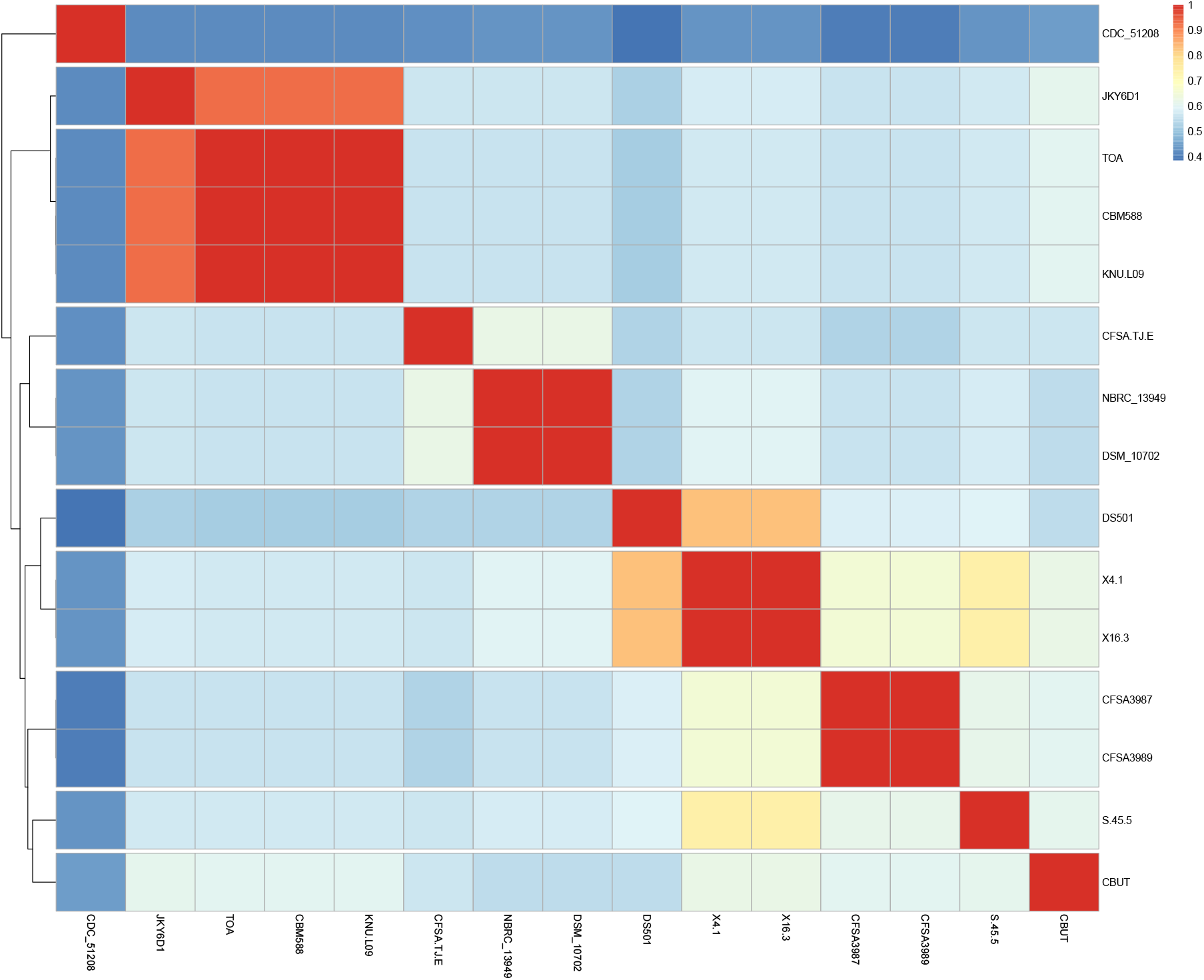
Phylogenetic tree showing relatedness of CBM588 with other strains of *C. butyricum*, with CDC_51208 used as the reference strain. Colours in the heatmap correspond to similarity; dark blue being most dissimilar and dark red most similar.

### 3.3 Characterisation of the CRISPR-Cas system in CBM588

The use of endogenous CRISPR for genetic engineering is an attractive alternative to the widely used CRISPR-Cas9 system from *Streptococcus pyogenes* as it removes the need for expressing the highly toxic Cas9 nuclease which results in low efficiency transformation. The endogenous CRISPR-Cas system of *C. butyricum* strain KNU-L09 was previously deployed for genome editing and proof of concept also performed in CBM588 (46). By interrogating the CBM588 megaplasmid (accession CP132345) with the CRISPRCasFinder webtool (47), a functional CRISPRCas type I-B locus was identified. CRISPR arrays contain spacers derived from previous invading organisms separated by direct repeats (DR). Homology between the spacers and new invading sequences facilitates the degradation of the invading element. A protospacer adjacent motif (PAM) at the 5’ end of the spacer allows the organism to distinguish between self and non-self sequences. Identification of the PAM is necessary for genome editing.

In CBM588, directly downstream from the CRISPR locus, an array was detected with 57 spacer sequences separated by the direct repeat (DR) GATTAACAGTAACATTAGATGTATTTAAAT. A large-scale analysis of CRISPR spacers by Vink *et al*., (2021) previously demonstrated a link between DR sequences and functional PAMs (48). Interestingly, the CBM588 DR sequence was identical to the DR of the CRISPR systems of 3 of the *C. butyricum* strains analysed in this study (KNU-L09, 29-1 and CDC_51208: accession **<u>CP013489</u>.<u>1</u>, <u>CP039703</u>.<u>1</u>** and **<u>CP013238</u>.<u>1</u>** respectively) and was found to be associated to a TCA PAM sequence (48). However, Zhou *et al*, (2021) reported the successful engineering of CBM588 using the PAM sequence ACA (46). Future engineering of CBM588 could deploy ACA or TCA as functional PAM sequences.

### 3.4 Butyrate production pathway in CBM588

The butyrate formation pathway has been well studied in *C. butyricum* (5) and the molecular pathway previously characterised in strain KNU-L09 (30). First pyruvate is oxidised to acetyl coenzyme A (acetyl-CoA) by pyruvate:ferredoxin (flavodoxin) oxidoreductase or by formate acetyltransferase. Knowledge of the enzymes required for this pathway in KNU-L09 enabled identification of the genes encoding these enzymes in CBM588. The molecular pathway from acetyl-CoA to butyrate with gene IDs is shown in Figure 2. All genes encoding the enzymes of this pathway were found exclusively on the chromosome of CBM588.

Other than *thlA* (CB_05870), the rest of the genes were organised as a cluster or as a pair. The following genes; butyryl-CoA dehydrogenase (*bcd*), 3-hydroxybutyryl-CoA dehydrogenase (*hbd*), crotonase (*crt*) and the EtfA/B complex were found on the cluster (Figure 3). A second *bcd* gene was found on the chromosome, CB_16190, with 69% protein identity over 258/375 amino acids to CB_04920. Since this gene was not part of the cluster, its expression is unlikely to be associated or co-ordinated with other enzymes in this cluster to make butyrate. The 2 genes involved in the final step of butyrate production (*ptb* and *buk*) were found as a pair and likewise, the 2 genes involved in acetate production, *pta* and *ack* (Figure 2). Clusters of genes involved in butyrate production have been reported before for *C. butyricum* (30) as well as for other bacteria (49, 50).

**Figure 2.**
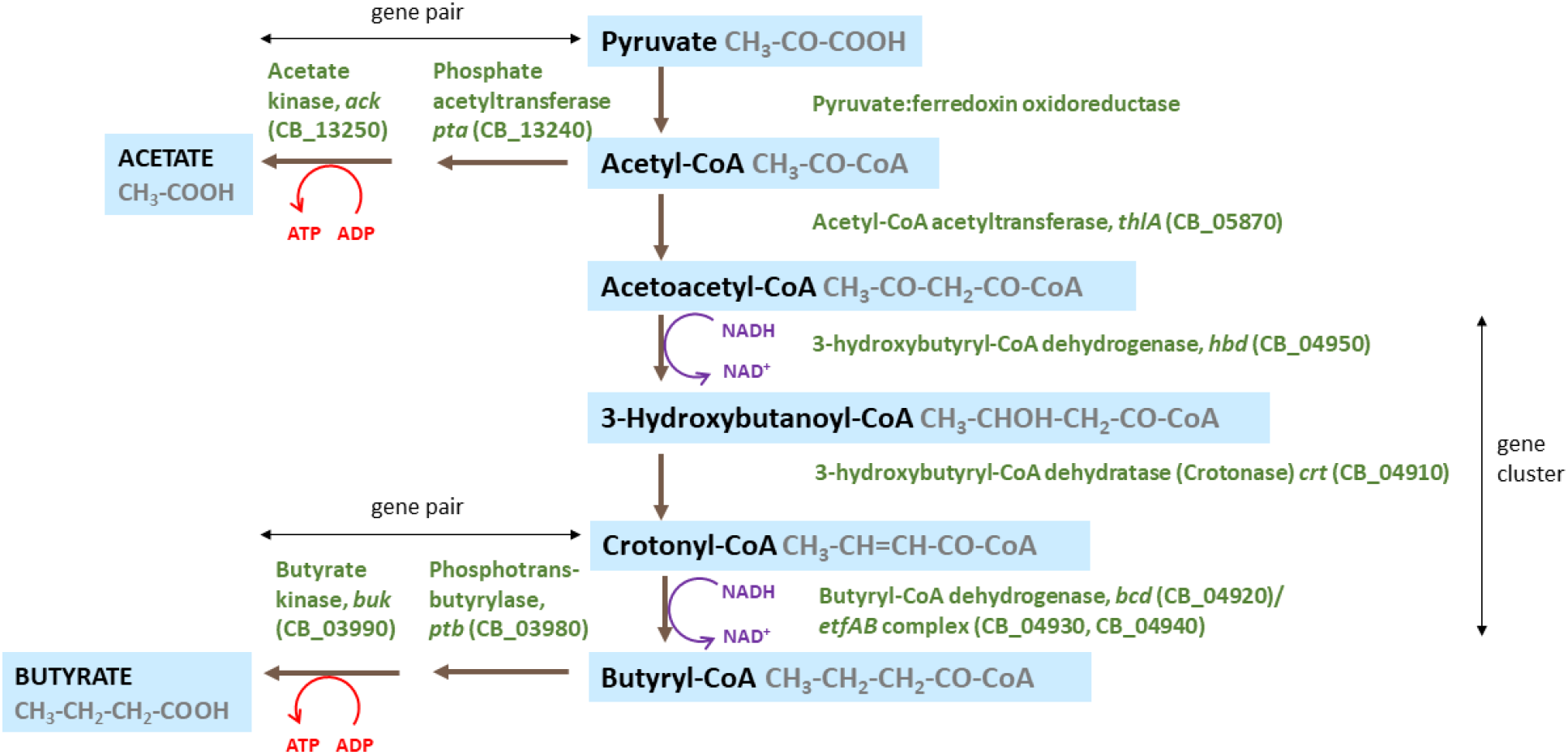
The molecular pathway for butyrate and acetate production pathway in *C. butyricum* strain CBM588.

#### 3.4.1 Formation of butyryl-CoA by electron-transferring flavoprotein (Etf) complex

The role of electron-transferring flavoprotein (Etf) complex in the butyrate pathway was previously investigated for *C. butyricum* strain KNU-L09 (30). The authors reported that after acetoacetyl-CoA is produced and reduced by Hbd, butyryl-CoA is formed following an endergonic ferredoxin reduction with NADH coupled to exergonic crotonyl-CoA reduction with NADH catalysed by the Bcd/EtfAB complex. The biochemistry of this reaction (2NADH+Fd_ox_ + crotonyl-CoA ➝ 2 NAD +Fd_red_ + butyryl-CoA) was first described for *Clostridium kluyveri* (51) then confirmed in other Firmicutes (52, 53). The identification of the same genes in the same clustered organisation on the chromosome (Figure 3) as seen in KNU-L09 suggests the above pathway for strain CBM588.

**Figure 3.**
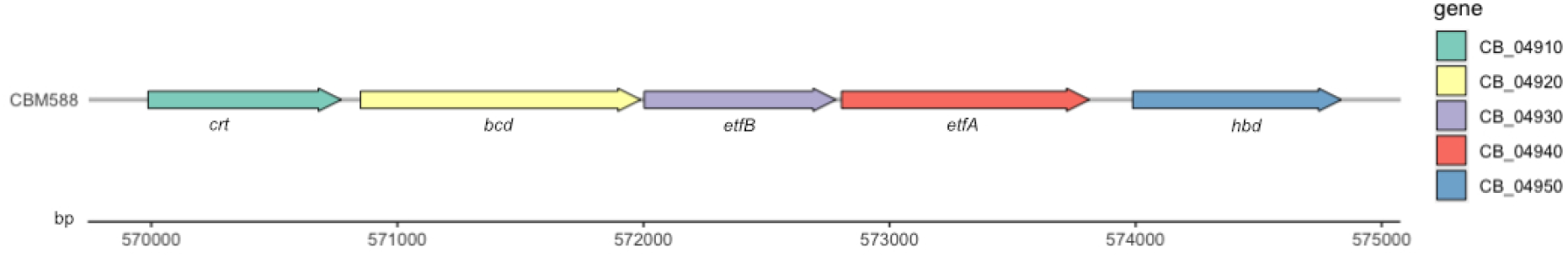
Clustered organisation of CB_04910 to CB_04950 on the chromosome of CBM588 involved in the conversion of acetoacetyl-CoA to butyryl-CoA.

#### 3.4.2 Final stage of butyrate pathway

The conversion of butyryl-CoA to butyrate in bacteria is catalysed either by butyryl-CoA:acetate CoA transferase (encoded by *but*) which transports the CoA component to external acetate resulting in the release of butyrate and acetyl-CoA, or by butyrate kinase (encoded by *buk*) after phosphorylation of butyryl-CoA by phosphotransbutyrylase encoded by *ptb*. To screen for butyrate producers, *but* and *buk* are common markers with the vast majority of strains possessing one or the other (5), however, some organisms possess both genes (5, 54). In the chromosome of CBM588, *buk* and *ptb* were present and *but* absent, as reported for all 24 strains of *C. butyricum* analysed by Zou *et al*., (2021) (55).

#### 3.4.3 Analysis of the cytosolic proteome to rank the abundance of enzymes involved in butyrate production

CBM588 was cultured to early stationary phase in *C. difficile* Minimal Medium (41). This medium lacks proteins resulting in less background contaminants for LC-MS analysis and contains glucose (10 g/L) important for butyrate production. Since the metabolic enzymes involved in butyrate production are cytoplasmic, total cytosolic proteins were harvested and LC-MS conducted to confirm the presence of these enzymes and to rank their abundance by relative MaxLFQ MS response. LFQ intensities are proven to be reliable proxies for relative protein abundance within complex samples (56, 57). A total of 1510 proteins were detected and all 8 enzymes in the butyrate production pathway found (Figure 2). Ptb was the least abundant enzyme (92^nd^) whilst ThlA was the most abundant (4^th^) (Figure 4) (Table A1). The Kruskal Wallis test was conducted to determine if there are any significant differences between any of the groups. Post-hoc Dunn’s Multiple Comparison tests were then performed and revealed a significant difference between Ptb and ThlA with p = 0.00769046. The levels of Buk and Crt were also low but not significantly lower than ThlA. CB_16190 encoded by the second *bcd* gene was not detected by LC-MS corroborating that its expression is not co-ordinated with the rest of the genes in the cluster (Figure 3) and unlikely to play a role in butyrate production.

**Figure 4.**
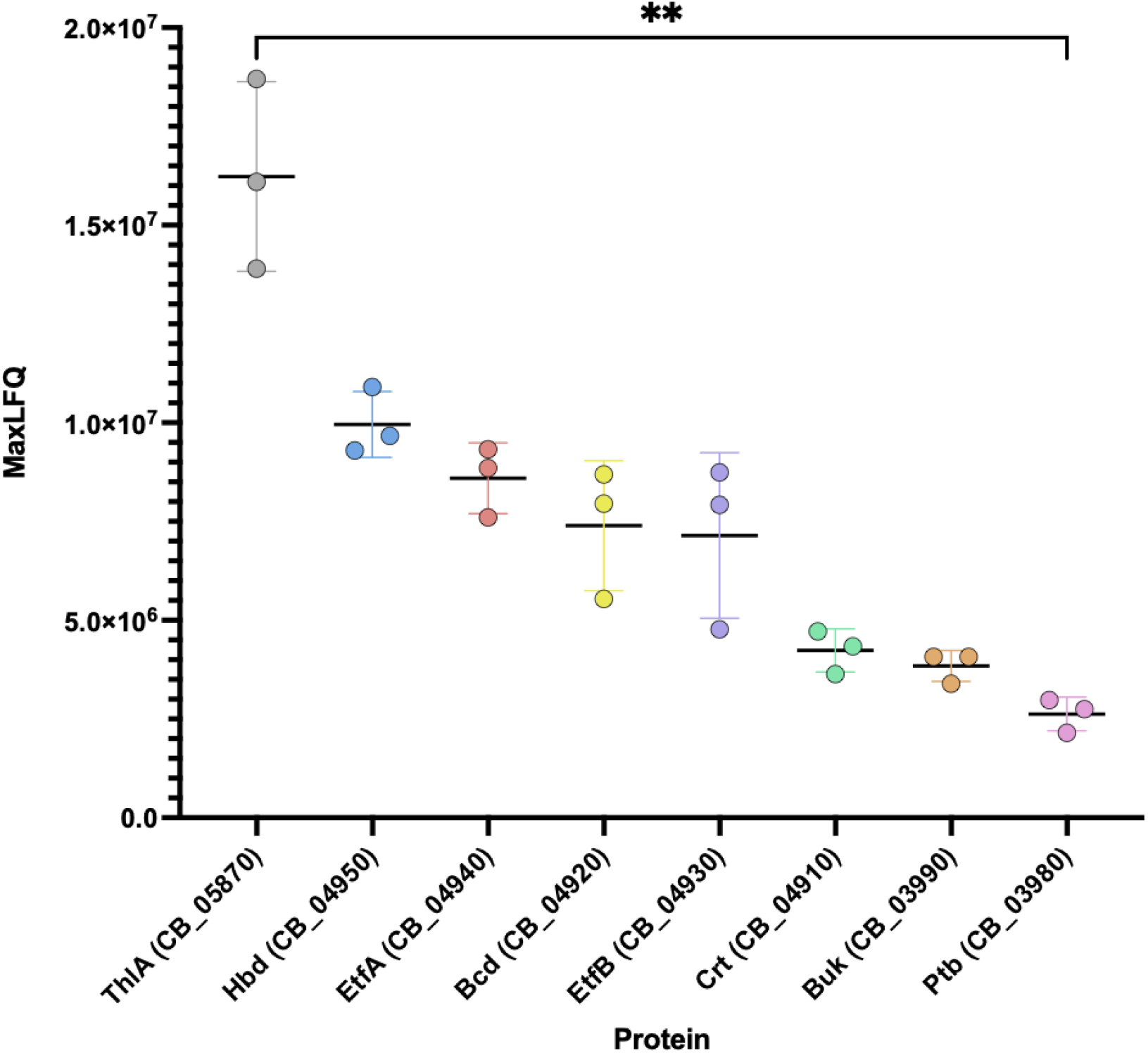
Dotplot of the relative MS intensities (MaxLFQ MS response) of enzymes involved in butyrate production analysed from total cytosolic proteins of 3 early stationary phase cultures of CBM588. Bars are medians, whiskers are inter-quartile-range and each dot is a single biological replicate (n=3). **Significant overall difference in MaxLFQ between enzymes by Kruskal Wallis test (p=0.00308) and specifically between ThlA and Ptb by Dunn’s post-hoc tests (adjusted for multiples comparisons, p=0.00769046).

### 3.5 High abundance proteins have an intact Shine-Dalgarno sequence while low abundance proteins contain SNPs

We set out to determine whether any associations could be made between the abundance of enzymes and regulatory sequences upstream of their coding sequences that determine their level of expression. Whilst gene editing tools for many species of *Clostridium* have greatly progressed over many years (58, 59), the characterisation of biological parts including promoters and ribosomal binding sites (RBSs) and terminators requires further advancement with each part optimised for each strain chosen for bioengineering purposes (60). Despite the limitations of available online tools, the conventional RBS Shine-Dalgarno (SD) sequence “AGGAGG” originally identified in *Escherichia coli* (61) was clearly visible within 16 nucleotides upstream of the ATG. For the 5 most abundant enzymes the SD sequence was intact. Conversely for the 3 least abundant enzymes, a single nucleotide polymorphism (SNP) was detected which may result in reduced translation efficiency of these genes (Table 1).

**Table 1.**
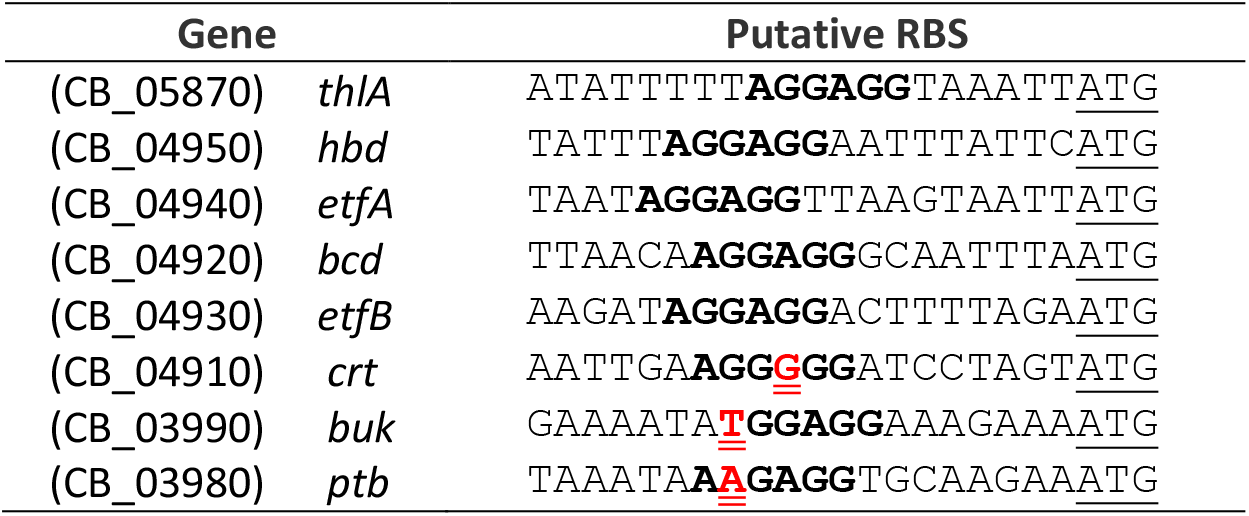
The putative ribosomal binding site (bold) upstream of the annotated translation start codon (underlined) of the genes involved in the butyrate production pathway in strain CBM588. SNPs are shown in red with double underline.

To test if the above trend was likely to be by chance given the small number of enzymes analysed, we expanded our investigation by comparing the most abundant proteins with the least abundant proteins in our dataset of 1510 cytosolic proteins. Of the 40 most abundant proteins, 26 (65%) had an intact SD sequence compared to 6 (15%) for the 40 least abundant (Table A2 and A3). By Mann– Whitney U/Wilcoxon rank-sum test, the difference in abundance (sum of 3 MaxLFQ intensity readings) between proteins encoded by genes with intact and non-intact SD sequences is highly statistically significant (p=0.0000387), suggesting a very strong association between intact SD sequences and high abundance of proteins (Figure 5).

**Figure 5.**
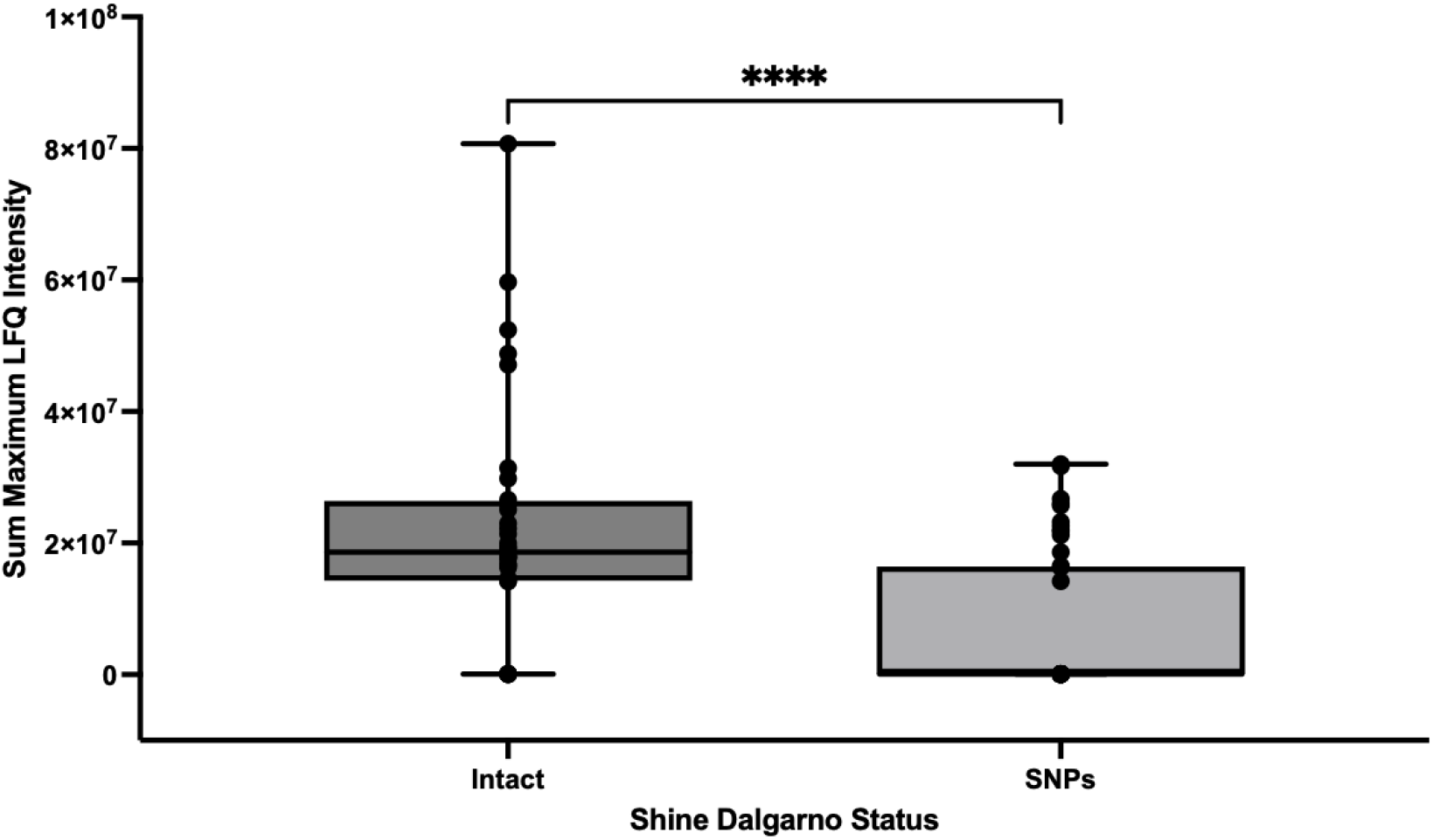
Boxplot showing abundance for the highest 40 and lowest 40 proteins and whether Shine-Dalgarno (SD) sequence is intact or contains SNPs. An intact SD correlates with higher protein abundance (measured as the sum of 3 MaxLFQ Intensity readings) compared to proteins where the gene encoding them has SNPs (single nucleotide polymorphisms) in the SD sequence. The difference in protein abundance for intact SD vs non-intact SD is highly significant by Mann–Whitney U/Wilcoxon rank-sum test (p=0.0000387). Lines are median, boxes are inter-quartile range and whiskers are minimum and maximum.

## 4 Discussion

Gut dysbiosis is associated with metabolic, gastrointestinal and neurological disorders as well as cancer (62). Short chain fatty acids (SCFA) produced by members of the colonic microbiota confer gut health by modulating intestinal immune homeostasis, improving intestinal barrier function and alleviating inflammation. Although the specific mechanisms accounting for the multitude of health benefits provided by *C. butyricum* are not fully elucidated, current evidence suggests that butyrate production in the colon is key. CBM588 is widely used in Japan, China and Korea (22) and has been authorised under the regulation of the European Parliament and of the Council as a novel food ingredient (63). Engineering this strain for enhanced butyrate production could render this probiotic all the more effective.

In this study we show that a combined genome-proteome approach was a facile method for identifying the rate-limiting enzymes of the butyrate production pathway in CBM588 based on abundance. First, whole genome sequencing was conducted using a combination of Illumina and PacBio. Next, the cytosolic proteome was investigated by LC-MS and relative MaxLFQ intensities measured to rank protein abundance (56, 57). By interrogating the genome to identify homologues of genes known to be involved in butyrate production in strain KNU-L09 (30) and comparing the relative abundance of the corresponding enzymes in CBM588, Ptb was found to be the least abundant followed by Buk and Crt.

Analysis of regulatory sequences upstream of the 8 genes involved in the butyrate production pathway revealed an intact Shine-Dalgarno (SD) sequence for all genes other than *ptb, buk* and *crt* which contained SNPs. We speculated that the translation efficiency of these 3 genes may account for their reduced abundance. To further our investigation, the SD sequences of the 40 most abundant proteins and the 40 least abundant proteins in our LC-MS dataset for cytosolic proteins were compared. The association of an intact SD with strong abundance was highly significant. The importance of SD sequences in translation efficiency in *Clostridium* is not widely recognised and more studies investigating the association between intact SD sequences and gene expression levels/abundance are required in this organism. One recent study investigating the transcriptome of *Clostridioides difficile* strain 630 identified the RBS to be aGGAGg for the majority of genes for which transcripts could be detected (64) suggesting high conservation of this sequence per se in this species. A similar finding was made for *Clostridium autoethanogenum* (65). Prior to the development of omic tools, studies focusing on individual genes have also reported AGGAGG as the likely RBS for other species of *Clostridium* including the type E toxin gene of *C. botulinum* (66), *colA* encoding collagenase of *C. perfringens* (67), *celA* gene encoding endoglucanase A of *C. themocellum* (68) to name but a few. Further investigations correlating expression level from transcriptomic and proteomic datasets with putative promoters, regulatory elements, RBSs and terminators under different culture conditions will not only improve our understanding of the biological parts required for strong gene expression in different species of *Clostridium* but expand our tool kit for bioengineering purposes.

Most engineering in *Clostridium* to elevate butyrate levels for product extraction has been performed in *C. tyrobutyricum*. Knock outs of *pta* and *ack* in the acetic acid biosynthesis pathway (Figure 2) resulted in mutants with a ∼14% decrease in acetic acid production and ∼30% higher butyrate production in fermentation (31, 33). Overexpression of *cat1* and *crt* enhanced the flux from acetyl-CoA to butyrate and significantly reduced acetic acid production, which resulted in a 2.24-fold increase in the butyric acid to acetic acid ratio (32). Further over-expression of phophofructokinase (*pfkA*) and pyruvate kinase (*pykA*) involved in the production of pyruvate resulted in simultaneously increased butyrate/acetate ratio, butyric acid concentration and productivity compared to the wild-type strain in batch fermentation using a high glucose concentration (32). Potential limitations posed by *C*.*tyrobutyricum* however is that this species has a narrow substrate spectrum and can use only a few monosaccharides (glucose, xylose and fructose), mannitol and lactate for growth (11). This can be attributed to its small genome which lacks genes for transport and catabolism of starch and disaccharides such as maltose and sucrose (11). In contrast, *C. butyricum* has a larger genome and can produce butyric acid from a wide variety of carbon sources including starch, disaccharides and glucose (18). Work conducted in *C. butyricum* to elevate butyrate levels has focused on inhibiting the transformation of acetyl-CoA to ethanol (rather than blocking acetate production) by deleting *adhE* and *aldh* encoding bifunctional aldehyde/alcohol dehydrogenase and aldehyde dehydrogenase respectively. The *adhE* and *aldh* mutants of strain RH-2 resulted in the increase in butyrate production by 61.9% and 49.5%, respectively, and meanwhile the decrease in ethanol production (46).

Applying the new knowledge generated by our study with what is already known for *C. tyrobutyricum* and *C. butyricum*, the proposed strategy for creating an enhanced probiotic would be to delete *ack* and *pta* to block acetate formation (Figure 2) and *adhE* and *aldh* to block ethanol formation and drive the carbon flux from acetyl-CoA to butyrate, and additionally over-express *ptb, buk* and *crt* in the chromosome. The RBS, promoter and terminator of these genes can be replaced with more effective heterologous counterparts and the endogenous CRISPR-Cas systems exploited as shown for *C. butyricum* strain KNU-L09 (46) using TCA or ACA as functional PAM sequences.

*C. butyricum* engineered for enhanced butyrate production using plasmids to over-express the required genes as conducted in *C. tyrobutyricum* could also be investigated. The impact of knocking out *pta* and *ack* in *C. butyricum* may have less impact on growth than found for *C. tyrobutyricum* (32) given its larger genome that provides more capabilities. It is important to take into account external factors known to impact upon butyric acid fermentation. The medium pH significantly affects cell growth and the final butyric acid/acetic acid ratio. Generally, a pH above 6.0 is desirable for butyric acid production, and pH6.3 is optimal for *C. tyrobutyricum* and may be applicable for *C. butyricum* (31). Another important factor to consider is that butyrate production in fermentation is typically inhibited by butyric acid (69, 70). Specifically, its accumulation in cells leads to a decrease in the intracellular pH as well as the ATP available for cell growth (71). Like, *C. tyrobutyricum, C. butyricum* has natural ability to tolerate high concentrations of acidic product however cell tolerance to butyrate inhibition may need to be improved by further engineering the strain and/or continuously removing butyric acid from the fermentation broth (extractive fermentation) (16, 70, 72, 73).

## 5 Conclusion

Studies identifying rate-limiting steps in metabolic pathways typically focus on establishing the activity of individual enzymes based on the inhibition or over-expression of the presumed rate-limiting step but can often be unsuccessful (74). What can be over-looked is the level of expression of enzymes. Those that are produced in low quantities quickly become saturated (regardless of their strength of activity) preventing further production of the end-product. Our study demonstrates a simple approach for establishing enzymes that are rate-limiting due to low abundance. As sequence technologies advance for rapid generation of long reads, such as Nanopore sequencing, more and more genomes are becoming available or can be generated quickly *de novo*. In parallel, continuous improvements are being made in LC-MS approaches such as increased LC-MS acquisition speeds (44, 75) and more readily available free-to-use software for data analysis (42, 43). This means that these two omics approaches can now be more easily adopted and combined to investigate rate-limiting steps in other important metabolic pathways and guide bioengineering to create strains with important industrial applications.

## Acknowledgements

We thank Dr Gareth Little for advice on sequencing approaches. The Illumina sequencing was conducted by DeepSeq at the University of Nottingham and PacBio at GenomeQuébec. We thank Jaime Hughes for administrative assistance with shipment of samples and Nigel Minton for use of equipment in the Nottingham BBSRC/EPSRC Synthetic Biology Research Centre.

The LC-MS work was conducted in the York Centre of Excellence in Mass Spectrometry (CoEMS) which was created thanks to a major capital investment through Science City York, supported by Yorkshire Forward with funds from the Northern Way Initiative, and subsequent support from EPSRC (EP/K039660/1; EP/M028127/1).

## Data availability statement

The genome of CBM588 has been deposited with the NCBI under GenBank accession numbers **<u>CP132344</u>, <u>CP132345</u>** and **<u>CP132346</u>**. The raw reads are available in the NCBI SRA under accession numbers **<u>SRX21335411</u>** and **<u>SRX21335410</u>** (BioProject number **<u>PRJNA1003819</u>**).

Raw mass spectrometry data and proteomic database results are deposited in MassIVE (MSV000092649, doi:10.25345/C5KW57V17) and referenced in ProteomeXchange (PXD044562).

## Appendix

**Table A1.** MaxLFQ Intensity of the cytosolic fraction of 3 early stationary phase cultures of strain CBM588.

| Enzyme | 1E_1 MaxLFQ Intensity | 2E_1 MaxLFQ Intensity | 3E_1 MaxLFQ Intensity | Sum MaxLFQ | Ranking position out of 1510 proteins |
| --- | --- | --- | --- | --- | --- |
| ThlA (CB_05870) | 1.61E+07 | 1.39E+07 | 1.87E+07 | 4.88E+07 | 4<sup>th</sup> |
| Hbd (CB_04950) | 9.67E+06 | 9.30E+06 | 1.09E+07 | 2.98E+07 | 9<sup>th</sup> |
| EtfA (CB_04940) | 8.85E+06 | 7.60E+06 | 9.33E+06 | 2.58E+07 | 13<sup>th</sup> |
| Bcd (CB_04920) | 7.95E+06 | 5.54E+06 | 8.69E+06 | 2.22E+07 | 19<sup>th</sup> |
| EtfB (CB_04930) | 7.92E+06 | 4.77E+06 | 8.74E+06 | 2.14E+07 | 22<sup>nd</sup> |
| Crt (CB_04910) | 4.34E+06 | 3.64E+06 | 4.72E+06 | 1.27E+07 | 47<sup>th</sup> |
| Buk (CB_03990) | 4.07E+06 | 3.39E+06 | 4.07E+06 | 1.15E+07 | 55<sup>th</sup> |
| Ptb (CB_03980) | 2.98E+06 | 2.15E+06 | 2.75E+06 | 7.88E+06 | 92<sup>nd</sup> |

**Table A2.**
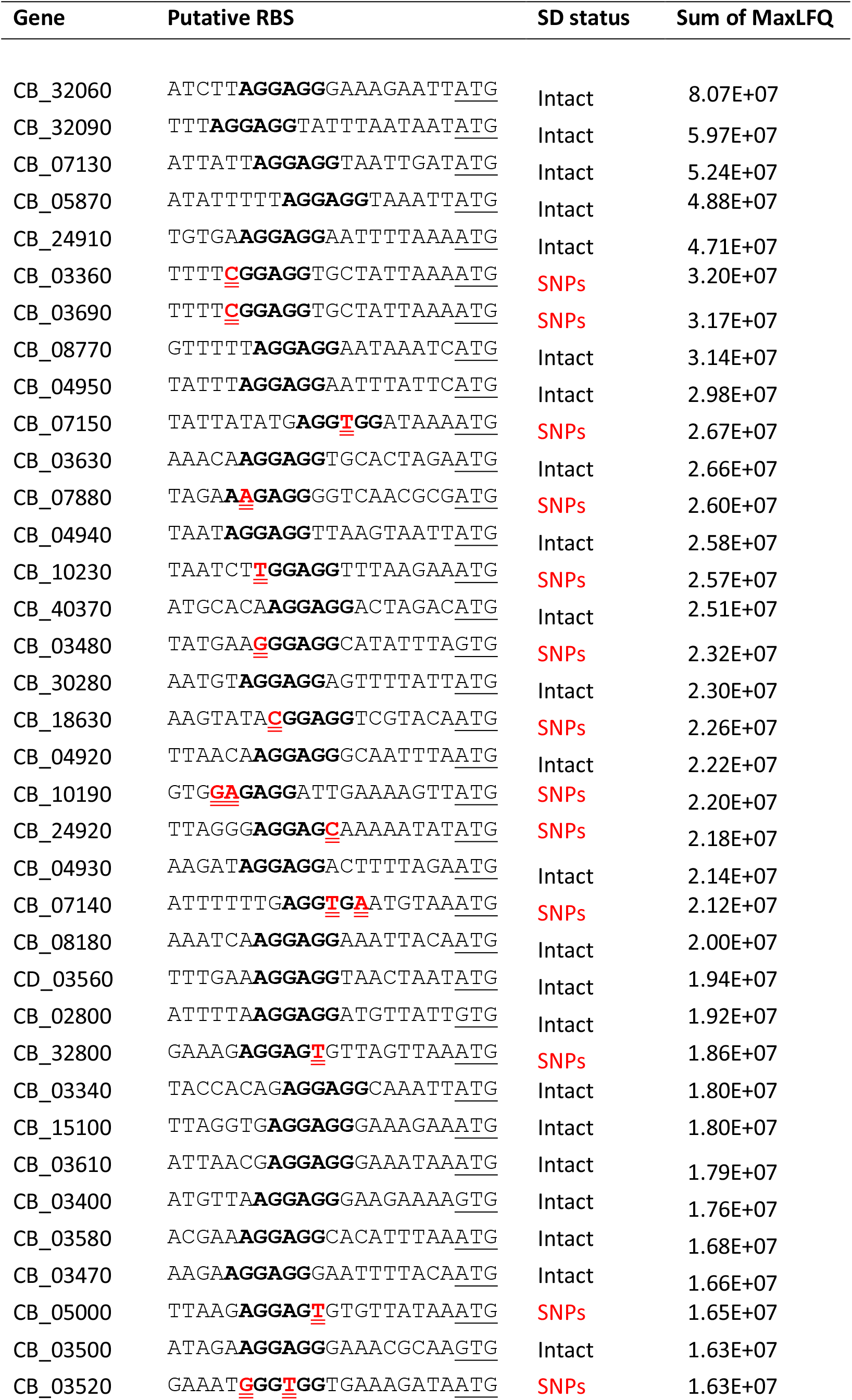
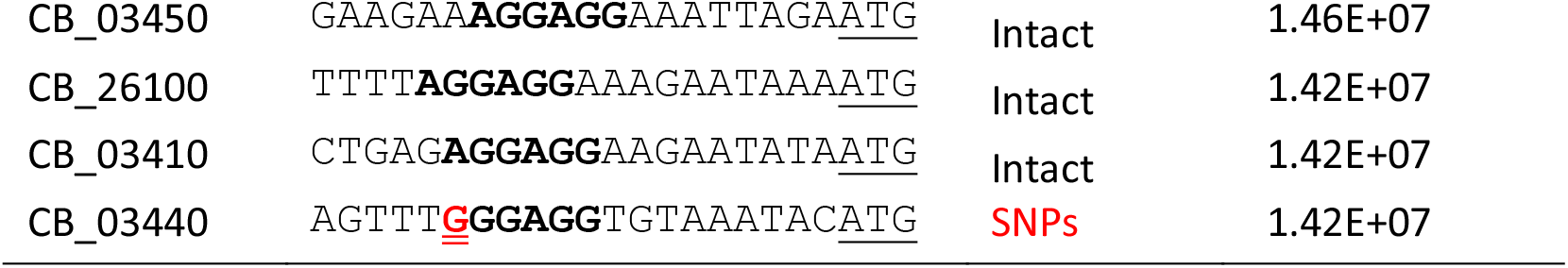
The highest abundant cytosolic proteins of strain CBM588 and their putative RBS. Shine-Dalgarno (SD) sequence (the consensus for which is AGGAGG) is shown in bold and SNPs in red with double underline. Annotated translation start codons are single underlined.

**Table A3.**
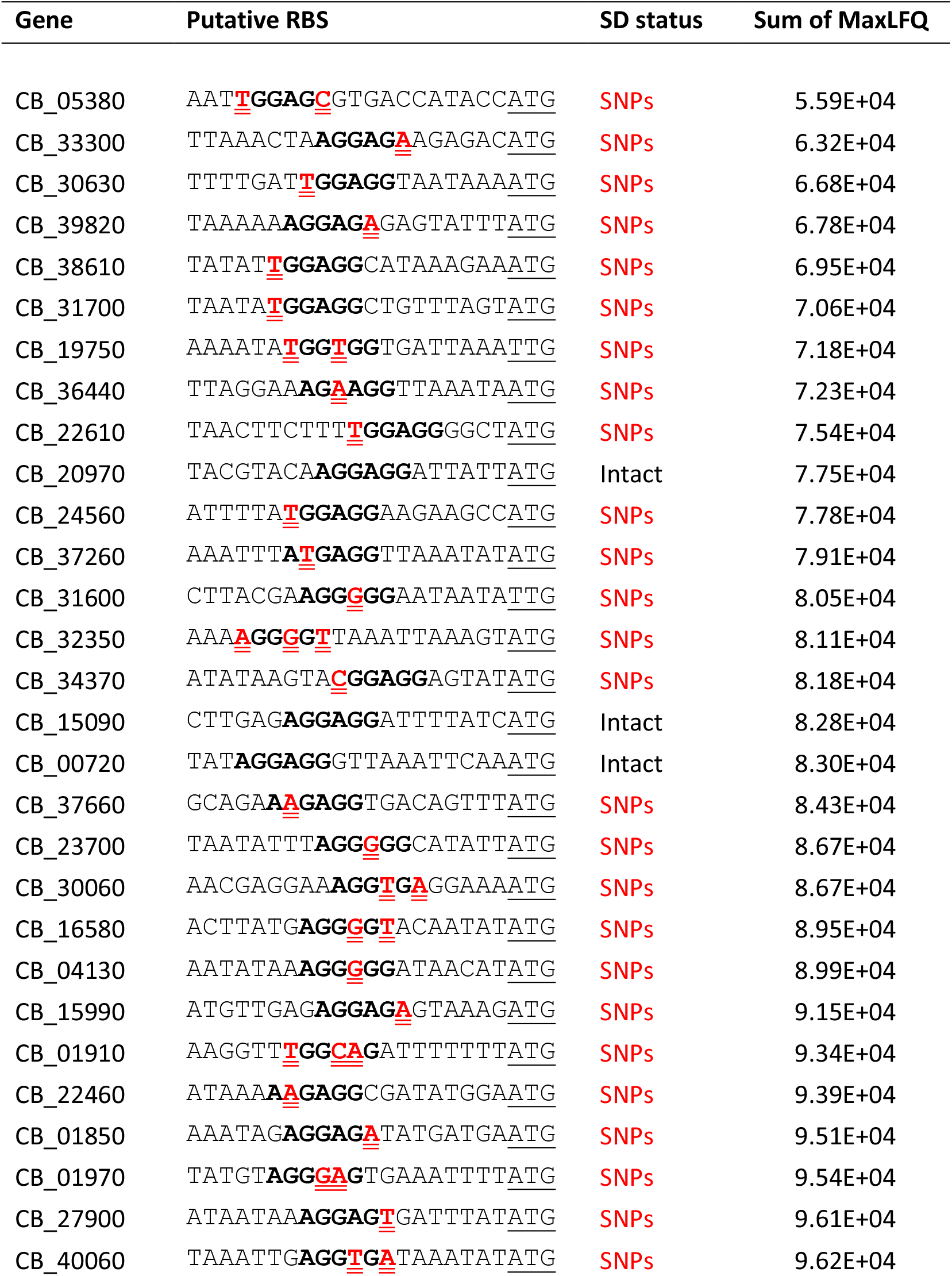
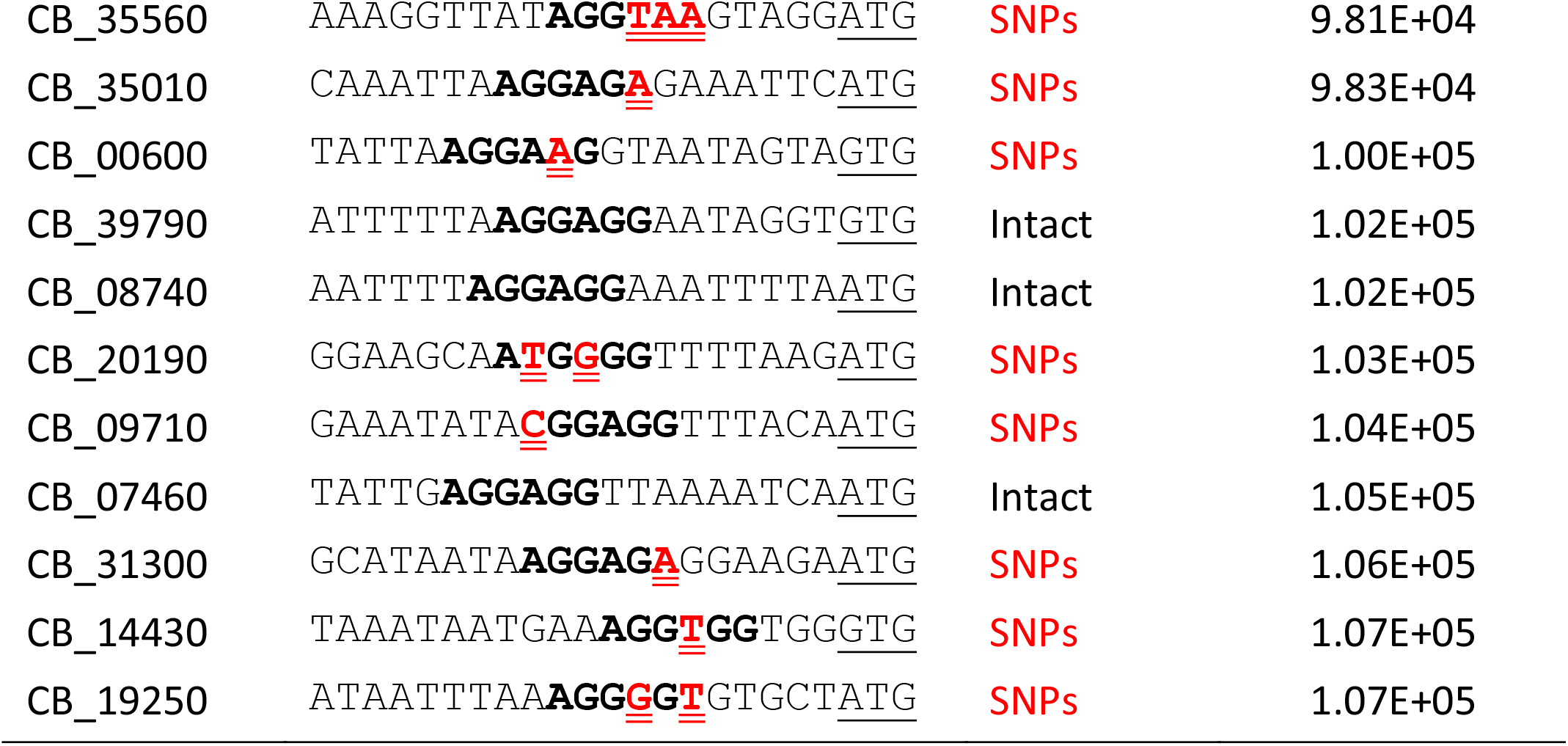
The lowest abundant cytosolic proteins of strain CBM588 and their putative RBS. Shine-Dalgarno (SD) sequence (the consensus for which is AGGAGG) is shown in bold and SNPs in red with double underline. Translation start codons are single underlined.

## References

1. Zhu LB, Zhang YC, Huang HH, Lin J. Prospects for clinical applications of butyrate-producing bacteria. World journal of clinical pediatrics. 2021;10(5):84–92.

2. Dwidar M, Park JY, Mitchell RJ, Sang BI. The future of butyric acid in industry. TheScientificWorldJournal. 2012;2012:471417.

3. Wang J, Lin M, Xu M, Yang ST. Anaerobic Fermentation for Production of Carboxylic Acids as Bulk Chemicals from Renewable Biomass. Advances in biochemical engineering/biotechnology. 2016;156:323–61.

4. Holscher HD. Dietary fiber and prebiotics and the gastrointestinal microbiota. Gut microbes. 2017;8(2):172–84.

5. Vital M, Howe AC, Tiedje JM. Revealing the bacterial butyrate synthesis pathways by analyzing (meta)genomic data. mBio. 2014;5(2):e00889.

6. Rivière A, Selak M, Lantin D, Leroy F, De Vuyst L. Bifidobacteria and Butyrate-Producing Colon Bacteria: Importance and Strategies for Their Stimulation in the Human Gut. Frontiers in microbiology. 2016;7:979.

7. Furusawa Y, Obata Y, Fukuda S, Endo TA, Nakato G, Takahashi D, et al. Commensal microbe-derived butyrate induces the differentiation of colonic regulatory T cells. Nature. 2013;504(7480):446–50.

8. Mortensen PB, Clausen MR. Short-chain fatty acids in the human colon: relation to gastrointestinal health and disease. Scandinavian journal of gastroenterology Supplement. 1996;216:132–48.

9. Celasco G, Moro L, Aiello C, Mangano K, Milasi A, Quattrocchi C, et al. Calcium butyrate: Anti-inflammatory effect on experimental colitis in rats and antitumor properties. Biomedical reports. 2014;2(4):559–63.

10. Encarnação JC, Abrantes AM, Pires AS, Botelho MF. Revisit dietary fiber on colorectal cancer: butyrate and its role on prevention and treatment. Cancer metastasis reviews. 2015;34(3):465–78.

11. Jiang L, Fu H, Yang HK, Xu W, Wang J, Yang ST. Butyric acid: Applications and recent advances in its bioproduction. Biotechnology advances. 2018;36(8):2101–17.

12. Playne M. Propionic and butyric acids. Comprehensive biotechnology. 1985;3:731–59.

13. Scandola M, Ceccorulli G, Pizzoli M. Miscibility of bacterial poly (3-hydroxybutyrate) with cellulose esters. Macromolecules. 1992;25(24):6441–6.

14. Posey-Dowty JD, Seo KS, Walker KR, Wilson AK. Carboxymethylcellulose acetate butyrate in water-based automotive paints. Surface Coatings International Part B: Coatings Transactions. 2002;85(3):203–8.

15. Zhang C, Yang H, Yang F, Ma Y. Current progress on butyric acid production by fermentation. Current microbiology. 2009;59(6):656–63.

16. Zigová J, Šturdík E. Advances in biotechnological production of butyric acid. Journal of Industrial Microbiology and Biotechnology. 2000;24(3):153–60.

17. Jiang L, Wang J, Liang S, Cai J, Xu Z, Cen P, et al. Enhanced butyric acid tolerance and bioproduction by Clostridium tyrobutyricum immobilized in a fibrous bed bioreactor. Biotechnology and bioengineering. 2011;108(1):31–40.

18. Bao T, Feng J, Jiang W, Fu H, Wang J, Yang ST. Recent advances in n-butanol and butyrate production using engineered Clostridium tyrobutyricum. World journal of microbiology & biotechnology. 2020;36(9):138.

19. Mountzouris KC, McCartney AL, Gibson GR. Intestinal microflora of human infants and current trends for its nutritional modulation. The British journal of nutrition. 2002;87(5):405–20.

20. Corfield AP. The Interaction of the Gut Microbiota with the Mucus Barrier in Health and Disease in Human. Microorganisms. 2018;6(3).

21. MIYARISAN PHARMACEUTICAL CO., LTD. [cited 2023 12 August]. Clostridium butyricum MIYAIRI strain]. Available from: http://www.miyarisan.com/english_index.htm.

22. Seki H, Shiohara M, Matsumura T, Miyagawa N, Tanaka M, Komiyama A, et al. Prevention of antibiotic-associated diarrhea in children by Clostridium butyricum MIYAIRI. Pediatrics international : official journal of the Japan Pediatric Society. 2003;45(1):86–90.

23. Guinan J, Wang S, Hazbun TR, Yadav H, Thangamani S. Antibiotic-induced decreases in the levels of microbial-derived short-chain fatty acids correlate with increased gastrointestinal colonization of Candida albicans. Scientific reports. 2019;9(1):8872.

24. Hagihara M, Yamashita R, Matsumoto A, Mori T, Kuroki Y, Kudo H, et al. The impact of Clostridium butyricum MIYAIRI 588 on the murine gut microbiome and colonic tissue. Anaerobe. 2018;54:8–18.

25. Kuroiwa T, Kobari K, Iwanaga M. [Inhibition of enteropathogens by Clostridium butyricum MIYAIRI 588]. Kansenshogaku zasshi The Journal of the Japanese Association for Infectious Diseases. 1990;64(3):257–63.

26. Takahashi M, Taguchi H, Yamaguchi H, Osaki T, Kamiya S. Studies of the effect of Clostridium butyricum on Helicobacter pylori in several test models including gnotobiotic mice. Journal of medical microbiology. 2000;49(7):635–42.

27. Takahashi M, Taguchi H, Yamaguchi H, Osaki T, Komatsu A, Kamiya S. The effect of probiotic treatment with Clostridium butyricum on enterohemorrhagic Escherichia coli O157:H7 infection in mice. FEMS immunology and medical microbiology. 2004;41(3):219–26.

28. Woo TDH, Oka K, Takahashi M, Hojo F, Osaki T, Hanawa T, et al. Inhibition of the cytotoxic effect of Clostridium difficile in vitro by Clostridium butyricum MIYAIRI 588 strain. Journal of medical microbiology. 2011;60(Pt 11):1617–25.

29. Zhou K, Ng W, Cortés-Peña Y, Wang X. Increasing metabolic pathway flux by using machine learning models. Current opinion in biotechnology. 2020;66:179–85.

30. Detman A, Mielecki D, Chojnacka A, Salamon A, Błaszczyk MK, Sikora A. Cell factories converting lactate and acetate to butyrate: Clostridium butyricum and microbial communities from dark fermentation bioreactors. Microbial cell factories. 2019;18(1):36.

31. Liu X, Yang S-T. Kinetics of butyric acid fermentation of glucose and xylose by Clostridium tyrobutyricum wild type and mutant. Process Biochemistry. 2006;41(4):801–8.

32. Suo Y, Ren M, Yang X, Liao Z, Fu H, Wang J. Metabolic engineering of Clostridium tyrobutyricum for enhanced butyric acid production with high butyrate/acetate ratio. Applied microbiology and biotechnology. 2018;102(10):4511–22.

33. Zhu Y, Liu X, Yang ST. Construction and characterization of pta gene-deleted mutant of Clostridium tyrobutyricum for enhanced butyric acid fermentation. Biotechnology and bioengineering. 2005;90(2):154–66.

34. Hughes J, Aston C, Kelly ML, Griffin R. Towards Development of a Non-Toxigenic Clostridioides difficile Oral Spore Vaccine against Toxigenic C. difficile. Pharmaceutics. 2022;14(5).

35. Kolmogorov M, Yuan J, Lin Y, Pevzner PA. Assembly of long, error-prone reads using repeat graphs. Nature biotechnology. 2019;37(5):540–6.

36. Camacho C, Coulouris G, Avagyan V, Ma N, Papadopoulos J, Bealer K, et al. BLAST+: architecture and applications. BMC bioinformatics. 2009;10:421.

37. Tanizawa Y, Fujisawa T, Nakamura Y. DFAST: a flexible prokaryotic genome annotation pipeline for faster genome publication. Bioinformatics (Oxford, England). 2018;34(6):1037–9.

38. Dykes JK, Lúquez C, Raphael BH, McCroskey L, Maslanka SE. Laboratory Investigation of the First Case of Botulism Caused by Clostridium butyricum Type E Toxin in the United States. Journal of clinical microbiology. 2015;53(10):3363–5.

39. Brown CT, Irber L. sourmash: a library for MinHash sketching of DNA. Journal of open source software. 2016;1(5):27.

40. Kolde R, Kolde MR. Package ‘pheatmap’. R package. 2015;1(7):790.

41. Cartman ST, Minton NP. A mariner-based transposon system for in vivo random mutagenesis of Clostridium difficile. Applied and environmental microbiology. 2010;76(4):1103–9.

42. Yu F, Haynes SE, Teo GC, Avtonomov DM, Polasky DA, Nesvizhskii AI. Fast Quantitative Analysis of timsTOF PASEF Data with MSFragger and IonQuant. Molecular & cellular proteomics : MCP. 2020;19(9):1575–85.

43. Demichev V, Messner CB, Vernardis SI, Lilley KS, Ralser M. DIA-NN: neural networks and interference correction enable deep proteome coverage in high throughput. Nature methods. 2020;17(1):41–4.

44. Meier F, Park MA, Mann M. Trapped Ion Mobility Spectrometry and Parallel Accumulation-Serial Fragmentation in Proteomics. Molecular & cellular proteomics : MCP. 2021;20:100138.

45. Nakanishi S, Tanaka M. Sequence analysis of a bacteriocinogenic plasmid of Clostridium butyricum and expression of the bacteriocin gene in Escherichia coli. Anaerobe. 2010;16(3):253–7.

46. Zhou X, Wang X, Luo H, Wang Y, Wang Y, Tu T, et al. Exploiting heterologous and endogenous CRISPR-Cas systems for genome editing in the probiotic Clostridium butyricum. Biotechnology and bioengineering. 2021;118(7):2448–59.

47. Couvin D, Bernheim A, Toffano-Nioche C, Touchon M, Michalik J, Néron B, et al. CRISPRCasFinder, an update of CRISRFinder, includes a portable version, enhanced performance and integrates search for Cas proteins. Nucleic acids research. 2018;46(W1):W246–w51.

48. Vink JNA, Baijens JHL, Brouns SJJ. PAM-repeat associations and spacer selection preferences in single and co-occurring CRISPR-Cas systems. Genome biology. 2021;22(1):281.

49. Anand S, Kaur H, Mande SS. Comparative In silico Analysis of Butyrate Production Pathways in Gut Commensals and Pathogens. Frontiers in microbiology. 2016;7:1945.

50. Boynton ZL, Bennet GN, Rudolph FB. Cloning, sequencing, and expression of clustered genes encoding beta-hydroxybutyryl-coenzyme A (CoA) dehydrogenase, crotonase, and butyryl-CoA dehydrogenase from Clostridium acetobutylicum ATCC 824. Journal of bacteriology. 1996;178(11):3015–24.

51. Li F, Hinderberger J, Seedorf H, Zhang J, Buckel W, Thauer RK. Coupled ferredoxin and crotonyl coenzyme A (CoA) reduction with NADH catalyzed by the butyryl-CoA dehydrogenase/Etf complex from Clostridium kluyveri. Journal of bacteriology. 2008;190(3):843–50.

52. Buckel W, Thauer RK. Flavin-Based Electron Bifurcation, A New Mechanism of Biological Energy Coupling. Chemical reviews. 2018;118(7):3862–86.

53. Demmer JK, Pal Chowdhury N, Selmer T, Ermler U, Buckel W. The semiquinone swing in the bifurcating electron transferring flavoprotein/butyryl-CoA dehydrogenase complex from Clostridium difficile. Nature Communications. 2017;8(1):1577.

54. Zhang Y, Yu M, Yang ST. Effects of ptb knockout on butyric acid fermentation by Clostridium tyrobutyricum. Biotechnology progress. 2012;28(1):52–9.

55. Zou W, Ye G, Zhang K, Yang H, Yang J. Analysis of the core genome and pangenome of Clostridium butyricum. Genome. 2021;64(1):51–61.

56. Al Shweiki MR, Mönchgesang S, Majovsky P, Thieme D, Trutschel D, Hoehenwarter W. Assessment of Label-Free Quantification in Discovery Proteomics and Impact of Technological Factors and Natural Variability of Protein Abundance. Journal of proteome research. 2017;16(4):1410–24.

57. Silva JC, Gorenstein MV, Li GZ, Vissers JP, Geromanos SJ. Absolute quantification of proteins by LCMSE: a virtue of parallel MS acquisition. Molecular & cellular proteomics : MCP. 2006;5(1):144–56.

58. Minton NP, Ehsaan M, Humphreys CM, Little GT, Baker J, Henstra AM, et al. A roadmap for gene system development in Clostridium. Anaerobe. 2016;41:104–12.

59. Pyne ME, Bruder MR, Moo-Young M, Chung DA, Chou CP. Harnessing heterologous and endogenous CRISPR-Cas machineries for efficient markerless genome editing in Clostridium. Scientific reports. 2016;6:25666.

60. Joseph RC, Kim NM, Sandoval NR. Recent Developments of the Synthetic Biology Toolkit for Clostridium. Frontiers in microbiology. 2018;9:154.

61. Shine J, Dalgarno L. The 3’-terminal sequence of Escherichia coli 16S ribosomal RNA: complementarity to nonsense triplets and ribosome binding sites. Proceedings of the National Academy of Sciences of the United States of America. 1974;71(4):1342–6.

62. Stoeva MK, Garcia-So J, Justice N, Myers J, Tyagi S, Nemchek M, et al. Butyrate-producing human gut symbiont, Clostridium butyricum, and its role in health and disease. Gut microbes. 2021;13(1):1–28.

63. The European Commission. Commission Implementing Decision of 11 December 2014 authorising the placing on the market of Clostridium butyricum (CBM 588) as a novel food ingredient under Regulation (EC) No 258/97 of the European Parliament and of the Council (notified under document C (2014) 9345).. Official Journal of the European Union 2014.

64. Fuchs M, Lamm-Schmidt V, Sulzer J, Ponath F, Jenniches L, Kirk JA, et al. An RNA-centric global view of Clostridioides difficile reveals broad activity of Hfq in a clinically important gram-positive bacterium. Proceedings of the National Academy of Sciences of the United States of America. 2021;118(25).

65. de Souza Pinto Lemgruber R, Valgepea K, Gonzalez Garcia RA, de Bakker C, Palfreyman RW, Tappel R, et al. A TetR-Family Protein (CAETHG_0459) Activates Transcription From a New Promoter Motif Associated With Essential Genes for Autotrophic Growth in Acetogens. Frontiers in microbiology. 2019;10:2549.

66. Fujii N, Kimura K, Murakami T, Indoh T, Yashiki T, Tsuzuki K, et al. The nucleotide and deduced amino acid sequences of EcoRI fragment containing the 5’-terminal region of Clostridium botulinum type E toxin gene cloned from Mashike, Iwanai and Otaru strains. Microbiology and immunology. 1990;34(12):1041–7.

67. Matsushita O, Yoshihara K, Katayama S, Minami J, Okabe A. Purification and characterization of Clostridium perfringens 120-kilodalton collagenase and nucleotide sequence of the corresponding gene. Journal of bacteriology. 1994;176(1):149–56.

68. Béguin P, Cornet P, Aubert JP. Sequence of a cellulase gene of the thermophilic bacterium Clostridium thermocellum. Journal of bacteriology. 1985;162(1):102–5.

69. Peterson EC, Daugulis AJ. The use of high pressure CO2 -facilitated pH swings to enhance in situ product recovery of butyric acid in a two-phase partitioning bioreactor. Biotechnology and bioengineering. 2014;111(11):2183–91.

70. Wu Z, Yang ST. Extractive fermentation for butyric acid production from glucose by Clostridium tyrobutyricum. Biotechnology and bioengineering. 2003;82(1):93–102.

71. Zhang C, Yang H, Yang F, Ma Y. Current progress on butyric acid production by fermentation. Current microbiology. 2009;59:656–63.

72. Du J, McGraw A, Lorenz N, Beitle RR, Clausen EC, Hestekin JA. Continuous fermentation of Clostridium tyrobutyricum with partial cell recycle as a long-term strategy for butyric acid production. Energies. 2012;5(8):2835–48.

73. Peterson EC, Daugulis AJ. Demonstration of in situ product recovery of butyric acid via CO2-facilitated pH swings and medium development in two-phase partitioning bioreactors. Biotechnology and bioengineering. 2014;111(3):537–44.

74. Moreno-Sánchez R, Saavedra E, Rodríguez-Enríquez S, Olín-Sandoval V. Metabolic control analysis: a tool for designing strategies to manipulate metabolic pathways. Journal of biomedicine & biotechnology. 2008;2008:597913.

75. Heil LR, Damoc E, Arrey TN, Pashkova A, Denisov E, Petzoldt J, et al. Evaluating the performance of the Astral mass analyzer for quantitative proteomics using data independent acquisition. bioRxiv : the preprint server for biology. 2023.

